# Parallel Synapses with Transmission Nonlinearities Enhance Neuronal Classification Capacity

**DOI:** 10.1101/2024.07.01.601490

**Authors:** Yuru Song, Marcus K. Benna

**Author notes:** Correspondence (M.K.B.).

## Abstract

Cortical neurons often establish multiple synaptic contacts with the same postsynaptic neuron. To avoid functional redundancy of these parallel synapses, it is crucial that each synapse exhibits distinct computational properties. Here we model the current to the soma contributed by each synapse as a sigmoidal transmission function of its presynaptic input, with learnable parameters such as amplitude, slope, and threshold. We evaluate the classification capacity of a neuron equipped with such nonlinear parallel synapses, and show that with a small number of parallel synapses per axon, it substantially exceeds that of the Perceptron. Furthermore, the number of correctly classified data points can increase superlinearly as the number of presynaptic axons grows.

When training with an unrestricted number of parallel synapses, our model neuron can effectively implement an arbitrary aggregate transmission function for each axon, constrained only by monotonicity. Nevertheless, successful learning in the model neuron often requires only a small number of parallel synapses.

We also apply these parallel synapses in a feedforward neural network trained to classify MNIST images, and show that they can increase the test accuracy. This demonstrates that multiple nonlinear synapses per input axon can substantially enhance a neuron’s computational power.

## Introduction

Synaptic connections play a crucial role in information transmission within neural systems. There is mounting evidence to suggest that cortical neurons are frequently connected by more than a single synapse. Initial findings predominantly focused on the somatosensory cortex [1–7], particularly the barrel cortex. Subsequent research has indicated that multiple connections between the same pair of (pre- and postsynaptic) neurons, which we will refer to as parallel synapses here, also exist in other brain areas, such as the hippocampus [8, 9] and visual cortex [4]. The number of parallel synapses varies across different brain regions and research methodologies. For instance, in the rat barrel cortex, 4 to 6 synaptic contacts between layer 4 and layer 2/3 neuron pairs have been reported [7], while later in silico reconstructions have identified 5 to 25 parallel synapses in the rat somatosensory cortex [10].

Synapses not only transmit signals between neurons, but they also undergo plasticity. The observed types of plasticity include short-term plasticity [11], homeostatic plasticity [12–14] and long-term plasticity [15–17]. Even synapses from the same neuron can express different forms of plasticity [18–20]. Thus, when two neurons are connected by multiple parallel synapses, each of these synapses might have different synaptic strengths and signal transmission properties. However, the impact of these potential variations in synapse properties on the computational power of neural circuits remains to be explored.

Recent studies have begun to provide an understanding of the potential benefits of parallel synapses. In [21], a phenomenological model was developed to explain the formation of parallel synapses. One stochastic model [22] aims to explain the distribution of the number of synapses between two neurons. It considers the synapses between two neurons a result of structural plasticity interacting with neuronal activity. Prior research has also approached multi-synaptic connections through the framework of Bayesian learning, proposing that multiple synapses on the same dendrite optimally estimate the input distribution and facilitate faster learning [23]. However, faster learning does not necessarily imply an increased capacity for memory or task execution. In [24], parallel synapses are proposed to have independent temporal filters capable of processing time-dependent inputs, with filters adaptable through synaptic plasticity. The study demonstrates that multiple synaptic contacts with dendritic filters can triple the neuron’s memory capacity of spatio-temporal patterns.

Previous works have also explored the concept of multi-weight connections between pairs of neurons in machine learning contexts [25, 26]. In [25], the authors introduced a model featuring multi-weight connections between neuron pairs, with each weight representing synaptic strength through a specific type of neurotransmitter. Meanwhile, the neuron model from [26] assumes repeated axonal inputs to the postsynaptic dendritic trees. In both models, inputs from various axons interact through dendritic nonlinearity before reaching the soma.

In our work, we focus specifically on the functional benefits of parallel synapses in the absence of dendritic nonlinearities that combine different inputs, in order to understand the computational advantages they can convey without additional mechanisms that would further complicate our models and conflate our results with the known benefits of such mechanisms.

To elucidate the computational benefits of parallel synapses, we examine the memory capacity of a single neuron connected to presynaptic neurons through parallel synapses. Specifically, we consider each synapse to be parameterized by a nonlinear monotonic function, with parameters optimized to store a large number of presynaptic activity patterns. We systematically investigate the relationship between memory capacity and the number of parallel synapses from presynaptic axons. Our model demonstrates a large memory capacity that increases with the number of presynaptic axons. Even a moderate number of parallel synapses can substantially enhance the memory capacity of the postsynaptic neuron.

We also extend our model to a limiting case in which the number of parallel synapses is unbounded. Here, the aggregate synaptic transmission function (summarizing the combined effect of a set of parallel synapses) becomes essentially an arbitrary monotonic function. In this scenario, we again observe a large memory capacity that increases with respect to the number of presynaptic axons. Interestingly, despite the availability of an unlimited number of parallel synapses, the model only requires a small number of them to achieve this memory capacity.

Additionally, to test the model’s ability to generalize (to inputs not used for parameter optimization), we apply parallel synapses in a feedforward neural network and train the model to perform digit classification on the MNIST data set [27]. We show that parallel synapses can significantly increase the testing accuracy compared to a neural network without parallel synapses. This is the case even when both models have an equal number of parameters.

## Results

### Mathematical characterization of parallel synapses

We consider the memory capacity of a single neuron, which receives inputs from different presynaptic axons. As illustrated in Fig. 1a, each axon can establish several synaptic connections with the neuron’s dendrites. We assume that the input from a particular axon is the same for these parallel synapses, and that they contribute additively to the current into the soma, i.e., we neglect dendritic nonlinearities. If the synaptic transmission functions for these parallel synapses are identical, they merely serve to boost the overall dynamic range [28] of the postsynaptic current and thus could be deemed redundant. Therefore, we assume that the synaptic transmission functions can vary and be independently learned. In particular, if the synaptic transmission functions of the parallel synapses were linear, the overall synaptic transmission function of these synapses would also be merely a linear function (so that additional parallel synapses would not enlarge the class of input-output mappings that the neuron can implement). Thus we presuppose that each synaptic transmission function is nonlinear (Fig. 1b), and more specifically, we model it as a sigmoid function *h*_*i,j*_(*x*_*i*_) of the input activity with three parameters:

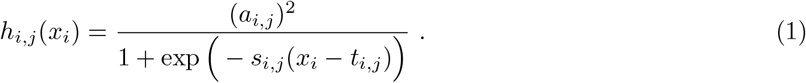

**Figure 1:**
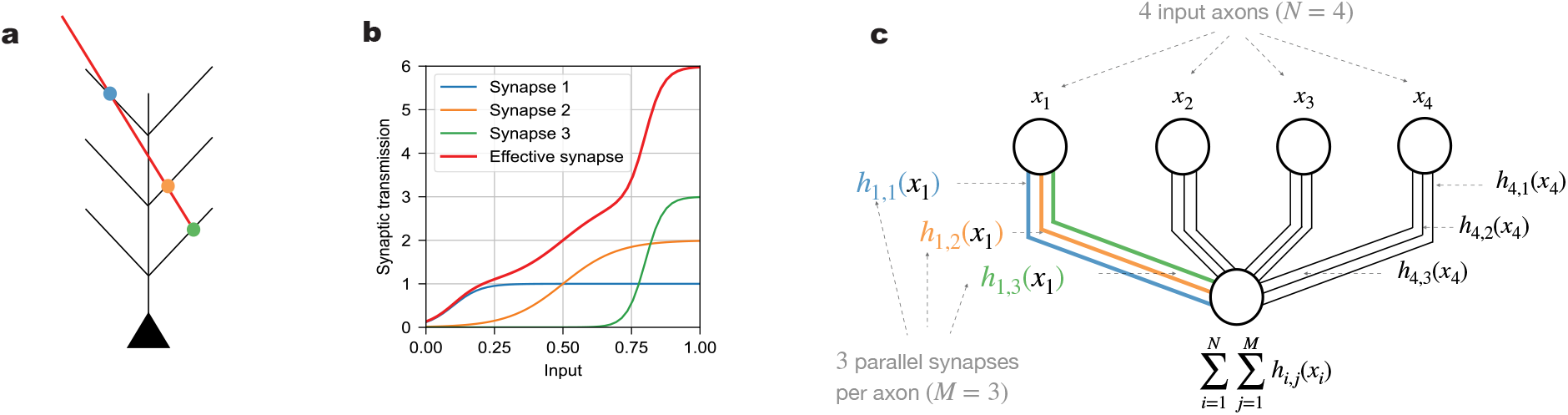
Neuronal connection via parallel synapses. **(a)**. Schematic of an axon (red) making multiple synaptic contacts with a postsynaptic neuron (black). The dots indicate three parallel synapses (blue, orange, and green). **(b)**. Each synapse has its own synaptic transmission function, parameterized as a sigmoid function. The effective, aggregate synaptic transmission function (red line) from the presynaptic axon to the postsynaptic neuron additively combines the parallel synapses’ transmission functions (blue, orange and green lines). **(c)**. Schematic of a neuron receiving inputs of presynaptic neuronal activity via parallel synapses. In the case shown, there are four axonal inputs (*N* = 4) and three parallel synapses (*M* = 3) per axon. Each presynaptic neuron (*i* = 1, 2, 3, 4) (upper four circles) connects to the postsynaptic neuron (lower circle) via three parallel synapses (*j* = 1, 2, 3), illustrated as a set of three parallel lines. The presynaptic neurons generally have different input activity values (*x*_1_, *x*_2_, *x*_3_ and *x*_4_), but the parallel synapses from a given presynaptic neuron convey the same input activity (e.g., the three synapses on the left have the same input value *x*_1_ from the first presynaptic neuron). The axon in (a) with three parallel synapses can be viewed as the first presynaptic input in (c). The blue, orange and green dots in (a) would then correspond to the blue, orange and green lines in (c), with synaptic transmission functions *h*_1,1_(*x*_1_), *h*_1,2_(*x*_1_) and *h*_1,3_(*x*_1_), respectively. These three synaptic transmission functions also correspond to the blue, orange and green lines in (b). The effective, aggregate synaptic transmission function from the first presynaptic neuronal input is *h*_1,1_(*x*_1_) + *h*_1,2_(*x*_1_) + *h*_1,3_(*x*_1_), plotted as the red line in (b). Since the model has no dendritic nonlinearities, the total input to the soma of the postsynaptic neuron is the sum of the activations from all parallel synapses of all presynaptic neurons, i.e.,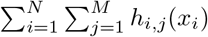.

In eqn. (1) and the following, we use *i* to index the *i*-th input axon, whose activity is denoted by *x*_*i*_, and *j* to index the *j*-th parallel synapse on that axon (Fig. 1c). The first parameter, (*a*_*i,j*_)^2^, is the amplitude of the transmission function, i.e., the maximum value of the synaptic response (which we write as a square to ensure it remains non-negative during learning). The second parameter, *t*_*i,j*_, is the threshold of the transmission function, which represents the value of presynaptic input at which the postsynaptic response increases most rapidly. The third parameter, *s*_*i,j*_, is proportional to the slope of the transmission function, which characterizes how steep it is at the threshold. Given that the response of excitatory synapses generally increases with larger input, we constrain the synaptic transmission function to increase monotonically (positive slope).

In contrast to models with nontrivial dendritic computation [29, 30], which are often functionally analogous to hidden layer neural networks that nonlinearly combine multiple inputs at an intermediate stage of processing, our model features nonlinearities that only act on individual input pathways (in addition to the final somatic output nonlinearity shared by all of these models). Thus our model has the same single-neuron architecture as the (standard) Perceptron [31] in which the structure of the dendritic tree of a real neuron plays no functional role, since the currents (into the soma) contributed by each synapse are assumed to simply add regardless of the location of the synapse. The only nonlinearity that integrates multiple inputs and facilitates their interaction is the nonlinear activation function at the soma, which implements the binary classification, modeling the neuron’s decision whether to spike or not in response to a certain input pattern.

### Parallel synapses can substantially increase the memory capacity of a neuron

To systematically quantify the memory capacity of our model neuron, we employ the classification of random patterns, a commonly used benchmark for measuring a model’s capacity [32]. In our case, successful classification is defined as correctly classifying all patterns in one dataset (Fig. 2a & b). The patterns to be classified are random; values from different input dimensions (i.e., presynaptic neuronal activities) are independently drawn from a uniform distribution between 0 and 1. The binary labels of these patterns are also random, sampled from a Bernoulli distribution with an equal probability of being either +1 or −1. For a given number of input axons *N*, the probability of successful classification decreases as the problem size *P* grows (Fig. 2c-e). Here we assume that the number of parallel synapses *M* is the same across all input axons. We want to determine the maximum problem size *P* a neuron can solve with 50% probability, denoted as *P*^∗^, normalized by the number of input axons *N*. We focus on the scaling behavior of the critical *P*^∗^*/N*, which describes the capacity of our model. For different *N* values, we test a set of problems with various sizes *P* to find the critical *P*^∗^. We repeat the process for different numbers of parallel synapses, i.e., values of *M* (Fig. 2c-e). The details of the gradient descent training algorithm used to optimize the model parameters and of the capacity estimation procedure for a given *M* and *N* are described in the Methods.

**Figure 2:**
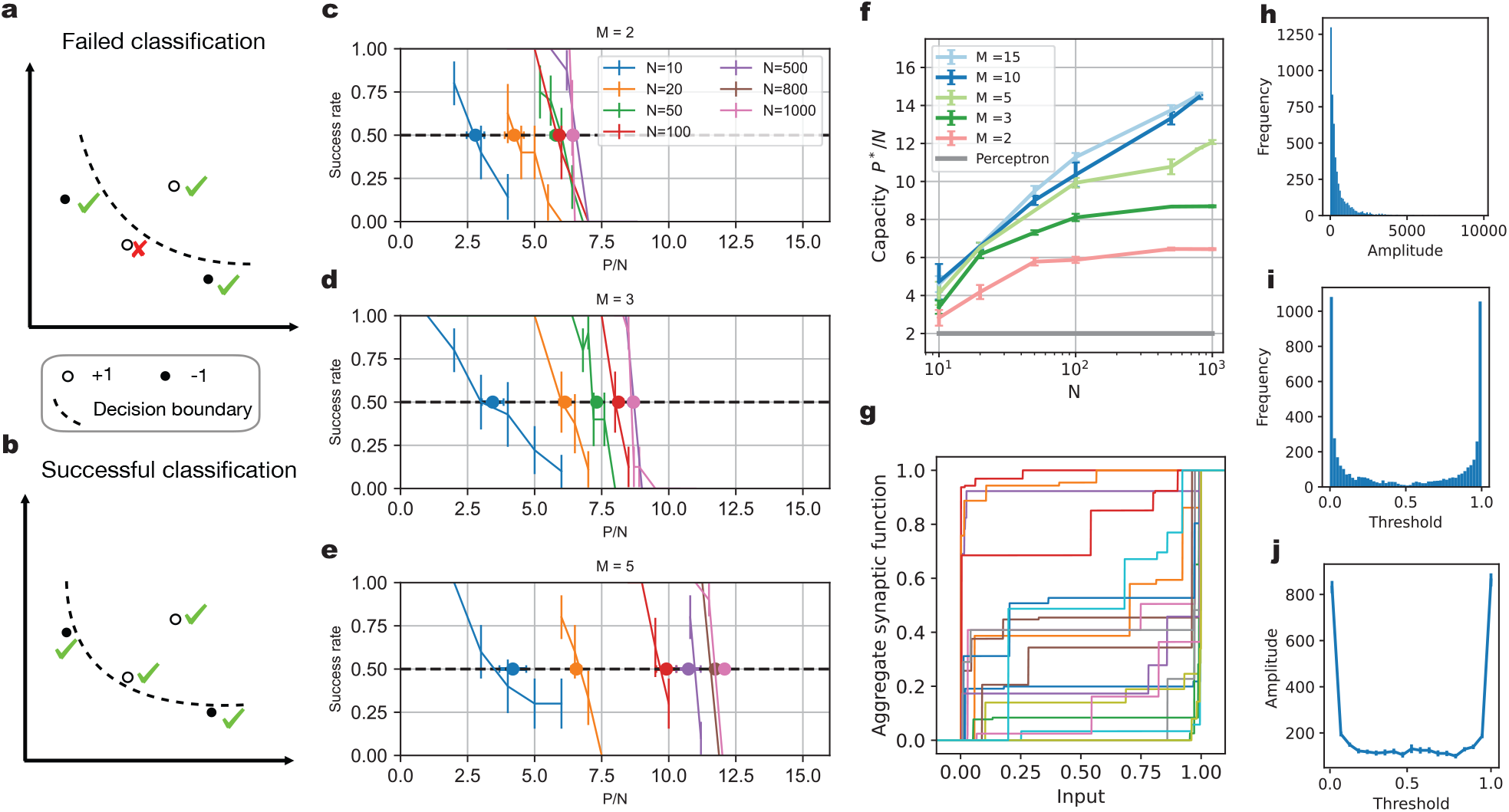
Increased memory capacity with parallel synapses. **(a-b)**. Illustration of binary classification task. In a two-dimensional input space, four random patterns are labeled with +1 (white dots) or −1 (black dots) labels. The goal is to find a decision boundary (dashed line) that correctly classifies all data points. In (a), the lower left data point with +1 label is misclassified, and thus the classification problem has not been solved yet. In (b), all data points are correctly classified, and thus the problem has been solved successfully. **(c-e)**. The success rate of correctly classifying a whole data set with *P* data points. For a neuron receiving *N* axons, each with *M* parallel synapses, the probability of completely solving the random classification tasks decreases as the number of patterns *P* grows. Results for *M* = 2, *M* = 3 and *M* = 5 are shown in panels (c), (d) and (e), respectively. We evaluate models with different numbers of input axons *N* (including 10, 20, 50, 100, 500, 800 and 1000), indicated by the color of the lines. **(f)**. The capacity *P*^∗^*/N* of a neuron with different numbers of input axons and different numbers of parallel synapses. With increasing *N*, the capacity keeps increasing, and thus *P*^∗^ grows faster than linearly. **(g)**. Examples of learned effective synaptic transmission functions, normalized by the maximum value. Here, *N* is 1000, *M* is 10 and *P* is 11000. **(h)**. Histogram of amplitude values for all synapses in (g). **(i)**. Histogram of threshold values for all synapses in (g). **(j)**. The relationship between parallel synapses’ amplitudes and their threshold values. The thresholds for all parallel synapses in (g) are divided into the same bins as in (i). We plot the average amplitude of the parallel synapses with thresholds in each bin.

Our comparison benchmark is the capacity of a neuron with only linear synaptic connections and a step-function somatic nonlinearity, the Perceptron model [31]. This model has a linear scaling of the critical *P*^∗^ with the number of presynaptic axons, with capacity *P*^∗^*/N* asymptotically equal to two for large *N* [33]. As shown in Fig. 2f, our model achieves a critical *P*^∗^*/N* substantially larger than two. Even with just two parallel synapses per axon (*M* = 2), the capacity *P*^∗^*/N* is about 6 with 100 different presynaptic neurons (*N* = 100). When the number of parallel synapses *M* increases, the capacity also grows. This shows that nonlinear parallel synapses can substantially enhance the classification capacity of a neuron. For a small number of parallel synapses per axons, such as *M* = 2, the capacity appears to saturate when the input axon number is large (Fig. 2f). With more parallel synapses, the capacity increases further and the saturation of the capacity seems to occur at larger *N* values.

Notably, the learned parallel synapses have a wide variety of response profiles, as illustrated by their aggregate synaptic transmission functions (Fig. 2g). For example, while most synapses have small amplitude, there are some synapses whose amplitudes are much larger (Fig. 2h). The synaptic function thresholds are more densely distributed near the edges of the input range (Fig. 2i). The amplitude of the synaptic response is also related to the synapse’s threshold (Fig. 2j). The synapses that have large amplitude tend to be sensitive to inputs near one of the edges of the input range, where their learned thresholds are preferentially located.

### Achieving high capacity only requires few synapses per axon

From the results of the previous section, it is clear that the presence of several nonlinear parallel synapses can increase the neuronal classification capacity. Does this mean that more parallel synapses are always better? Conversely, we can approach this from a theoretical perspective by asking: In the extreme case in which the number of parallel synapses on each input axon is unlimited, how large can the memory capacity become? To answer this question, we parameterize the effective aggregate transmission function of a set of parallel synapses as a staircase-like function of the axonal input (Fig. 5a). The height of each step can take an arbitrary non-negative value, and steps can appear at any value of the input. When the problem size is *P*, each aggregate transmission function for an axon can take at most *P* different values on this dataset, so with *P* steps we obtain a very flexible parameterization. For reasons of biological plausibility, we again constrain each aggregate transmission function to be monotonically increasing (by demanding non-negative step sizes). In order to clearly differentiate them, we will refer to the model with a limited number of parallel synapses on each input axon as a restricted neuron, and the model with an unlimited number of parallel synapses on each input axon as an unrestricted neuron. Again, we use binary classification problems with random patterns and random labels to assess the capacity of this system.

In the unrestricted neuron model, synaptic transmission functions involve step functions with discontinuities described by a large number of parameters. We have developed an alternative training algorithm for this model. This algorithm is based on a straightforward intuition: we use a loss function that increases the function values at input points with positive labels and decreases the function values for those with negative labels, pushing the model closer towards correct classification. This approach can be applied independently to each input dimension, and data points that remain misclassified can then be assigned increased importance for subsequent iterations. A detailed description of the algorithm is provided in the Methods.

Since there is no interaction between synaptic currents until the final somatic nonlinearity, we can construct a loss function for each input dimension (axon). We denote the value of the aggregate transmission function for the *µ*-th data point and the *i*-th axon by *I*_*i,µ*_, which depends on the input *x*_*i,µ*_. If the label *y*_*µ*_ of this data point is +1, our algorithm aims to ensure a sufficiently large value of the synaptic function, *I*_*i,µ*_ at *x*_*i,µ*_. On the other hand, if *y*_*µ*_ is −1, the algorithm adjusts *I*_*i,µ*_ to be small. To achieve this we initially define our cost function as 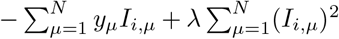, where we include a regularization term with parameter *λ* to discourage unrealistically large currents into the soma. Since some data points may be more challenging to classify than others, we introduce an importance weight *w*_*µ*_ for the loss of each data point. In summary, our loss function can be expressed as the following objective

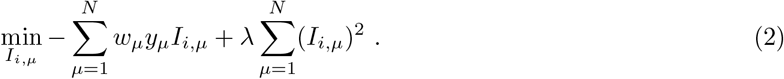

This single-axon objective can be used in an iterative algorithm to solve the classification problem. During each iteration, we first find a solution to eqn. (2) for all axons *i* (see Methods for details). Next, we evaluate the model’s predictions for the binary classification task. For misclassified data points, we increase their importance weights by one unit, i.e., 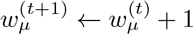. We then repeat this procedure until all data points are correctly classified or the maximum number of iterations is reached.

The unrestricted neuron model also has a memory capacity larger than that of the Perceptron (Fig. 3a). The scaling of the critical *P*^∗^*/N* value appears to increase with *N* (approximately in proportion to log *N*), rather than plateau when *N* is large. However, for the input dimensions (number of axons) relevant to realistic neural systems, say *N ∼* 1000, the unrestricted neuron has a capacity comparable to the restricted neuron model with *M* = 10 or *M* = 15. Interestingly, the unrestricted neuron does not utilize all available potential synapses (Fig. 3b & Fig. 3c). The majority of the utilized parallel synapses have thresholds near the edges of the input range (Fig. 3e). Also, the majority of the utilized parallel synapses have a small amplitude (Fig. 3d) and those parallel synapses with large amplitude are often tuned to the edges of the input range (Fig. 3b & Fig. 3f), in agreement with the restricted model, but with a bias towards the lower end of the input range (Fig. 3f). This bias arises from the regularization and the fact that we constrained the synaptic function to be nonnegative.

**Figure 3:**
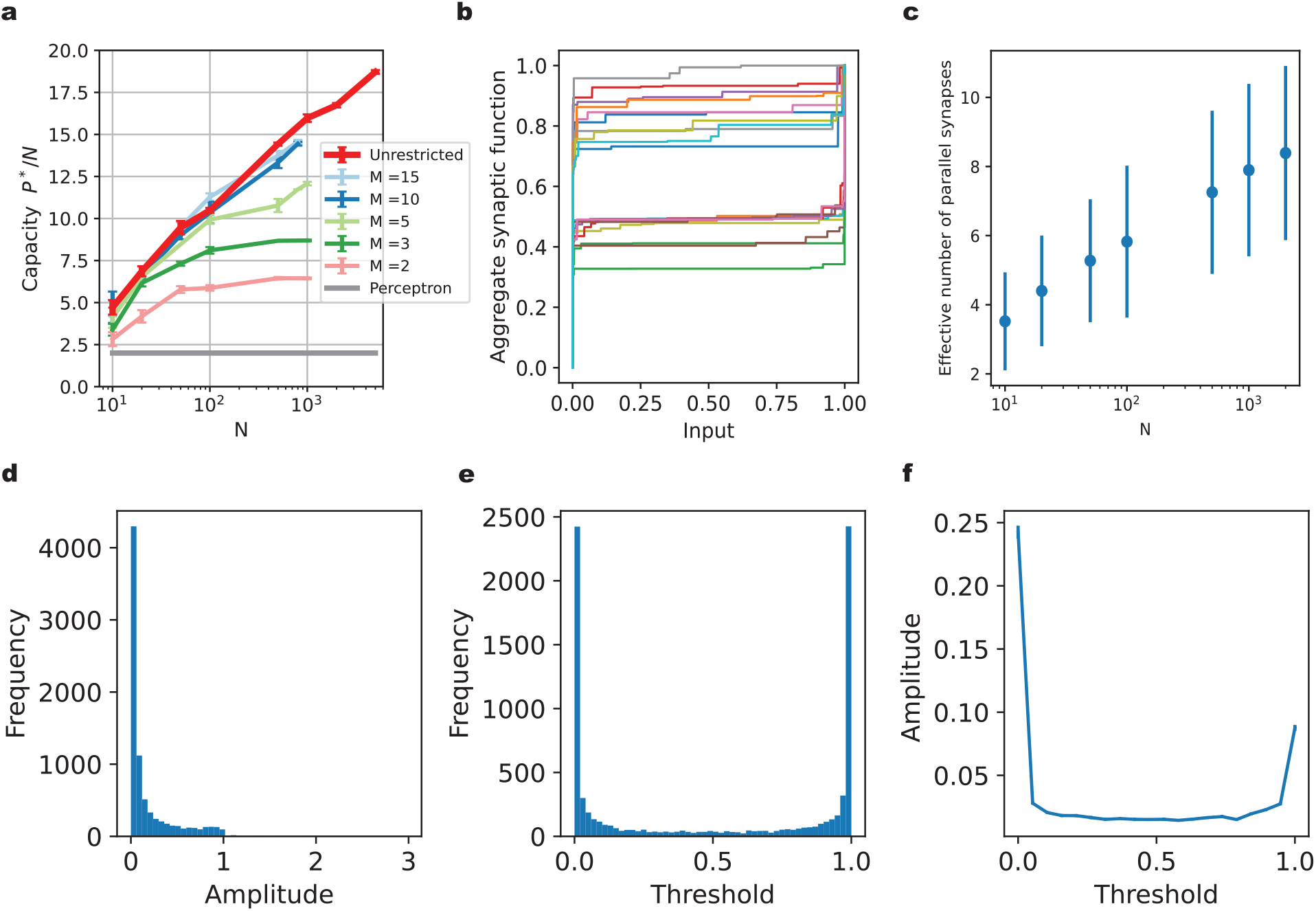
High memory capacity of the unrestricted model only requires few parallel synapses per axon. **(a)**. The capacity *P*^∗^*/N* of the unrestricted neuron (red line) compared to restricted neurons with different numbers of parallel synapses per axon. **(b)**. Examples of learned aggregate synaptic transmission functions for the unrestricted neuron, normalized by their maximum value. Here, *N* = 1000 and *P* = 16000. **(c)**.The effective number of parallel synapses on each input axon for the unrestricted neuron model, which varies with the number of input axons *N*. The problem size is near the critical number of patterns *P*^∗^. The effective number of parallel synapses is defined by counting the synapses with amplitude larger than 1/1000 of the maximum amplitude. **(d)**. Histogram of the amplitudes of all effective synapses from the example shown in (b) with *N* = 1000 and *P* = 16000. **(e)**. Histogram of the thresholds of all effective synapses from (b). **(f)**. Relationship between the amplitude of a synapse and its threshold. The thresholds for all effective parallel synapses in (b) are divided into the same bins as in (e). We plot the average amplitude of the synapses with thresholds in each bin.

### Enhanced classification accuracy through parallel synapses in neural networks

So far, we have used random patterns to evaluate the memory capacity of a single neuron equipped with parallel synapses. However, real-world problems often involve structured input distributions, and require generalization to unseen patterns. To evaluate the model’s ability to generalize, we incorporate parallel synapses into a feedforward neural network. We train this neural network to perform a classification task using the MNIST dataset, which consists of handwritten digits [27]. Specifically, we use a two-layer fully connected neural network, with variable number of neurons in the hidden layer (denoted as *D*_hidden_, between 5 and 30) and *D*_out_ = 10 neurons in the output layer, reflecting the one-hot encoding of the labels to be predicted. The input layer consists of *D*_in_ = 28 × 28 neurons, corresponding to the pixels of the images. Each neuron in the hidden layer connects to each output layer neuron via a fixed number (*M*) of parallel synapses (Fig. 4a). The connections between input layer and hidden layer are established through standard, linear synapses. We use rectified linear (ReLu) activation functions for neurons in both hidden layer and output layer. Parallel synapses are applied exclusively between the hidden and output layer neurons, adding *D*_hidden_*D*_out_(3*M* − 1) extra parameters compared to a network with the same architecture but only single, linear synapses. A two-layered neural network with parallel synapses has a total of (*D*_in_ + 1)*D*_hidden_ + (3*MD*_hidden_ + 1)*D*_out_ parameters, including the bias terms for the activities of each neuron in the hidden and output layers. For example, with *D*_hidden_ = 20 and *M* = 3 parallel synapses, the network has a total of 17510 parameters. For comparison, we also train a conventional two-layered neural network with *D*_hidden_ = 22 neurons in the hidden layer on the same MNIST task, using single, linear synapses for all neuronal connections (Fig. 4b). This network is referred to as ‘*D*_hidden_ = 22, linear synapse’ in Fig. 4c, and has roughly same parameter count as a network configuration with *D*_hidden_ = 20 and 3 parallel synapses, totaling 17500 learnable parameters. Similarly, for each of the *D*_hidden_ = 5, 10 and 30 cases, we train a corresponding network with linear synapses that has a roughly equal number of parameters. The total parameter count for both network types is detailed in the Supplementary Table 1. It’s important to note that because our parallel synapses have monotonically increasing transmission functions, i.e., are excitatory, we also apply this constraint to the linear synapses in the hidden to output layer weights of the standard (comparison) network, which thus also have to be nonnegative. We assess model performance by the testing accuracy on the held-out MNIST data.

**Figure 4:**
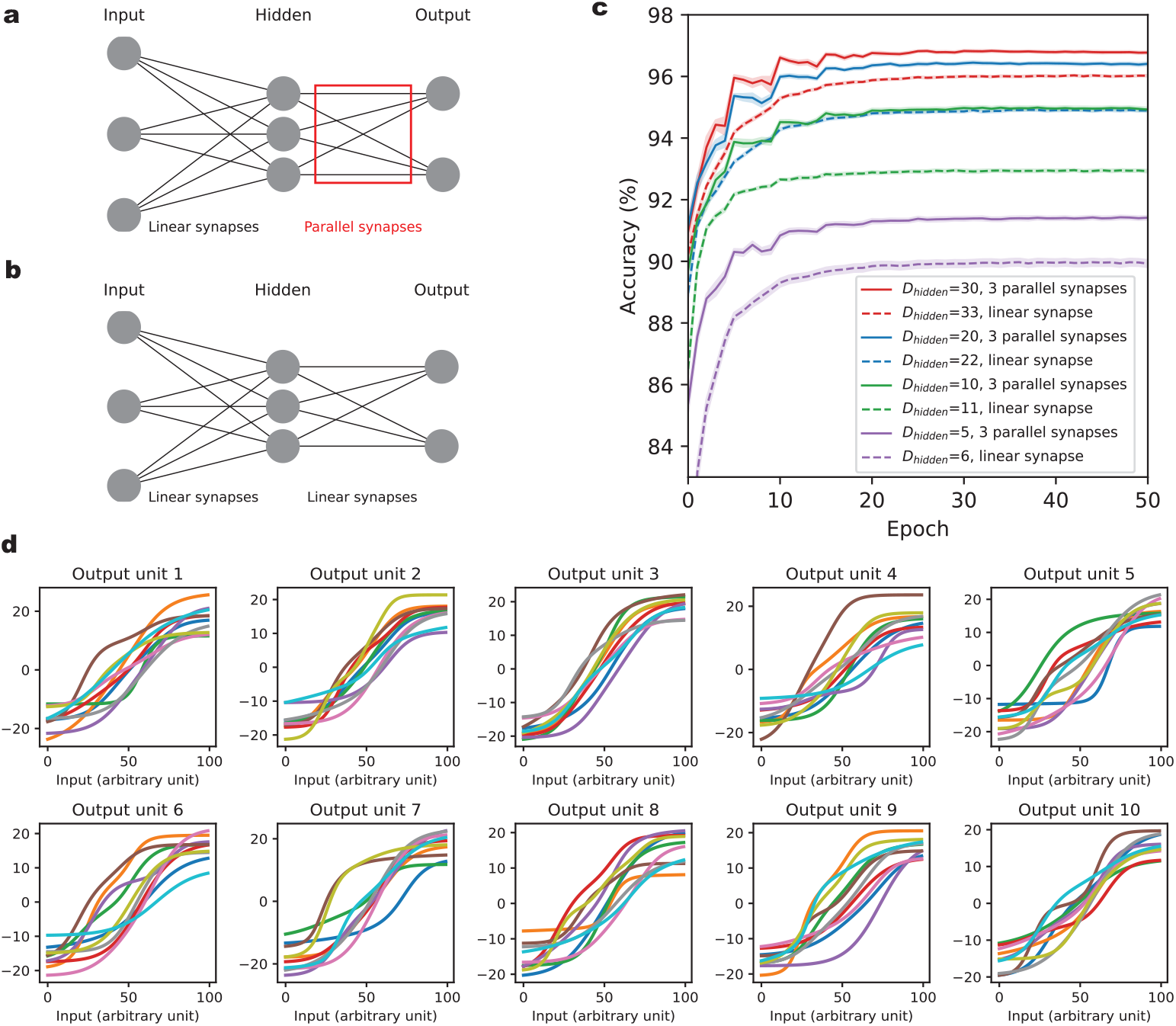
Enhancing classification accuracy for hand-written digits using parallel synapses. **(a)**. The feedforward network consists of an input layer, a hidden layer and an output layer. There are 784 and 10 neurons in the input and output layers, respectively, corresponding to the image pixels and the digit classes to be predicted. The number of neurons *D*_hidden_ in the hidden layer varies across different models. The connections between input and hidden layers are regular, linear synapses (no multiple parallel connections). The connections between hidden and output layers are nonlinear parallel synapses, with 3 parallel synapses per connection. **(b)**. The comparison benchmark model of a standard feedforward neural network with only linear synapses. There are 784 and 10 neurons in the input and output layers, respectively. The number of neurons in the hidden layer is slightly larger than in (a), in order to keep the total number of parameters in the two networks approximately the same. **(c)**. Classification accuracy on held-out data for neural network models with parallel synapses and standard neural networks with linear synapses for the MNIST data set. The x-axis indicates training epochs. The y-axis corresponds to the percentage of correctly classified test data point. Each model is initialized with 20 distinct random seeds, with results averaged across these seeds. **(d)**. Learned aggregate synaptic transmission functions for an example network with *D*_hidden_ = 10. For each pair of hidden neuron and output neuron, there are 3 parallel synapses connecting them. The aggregate synaptic transmission functions are grouped together according to the output neuron to which they connect, each corresponding to one panel. The x-axis is the percentile of input current to the corresponding hidden unit, as the parallel synapses in one panel share the same input to the hidden unit from which they originate.

**Figure 5:**
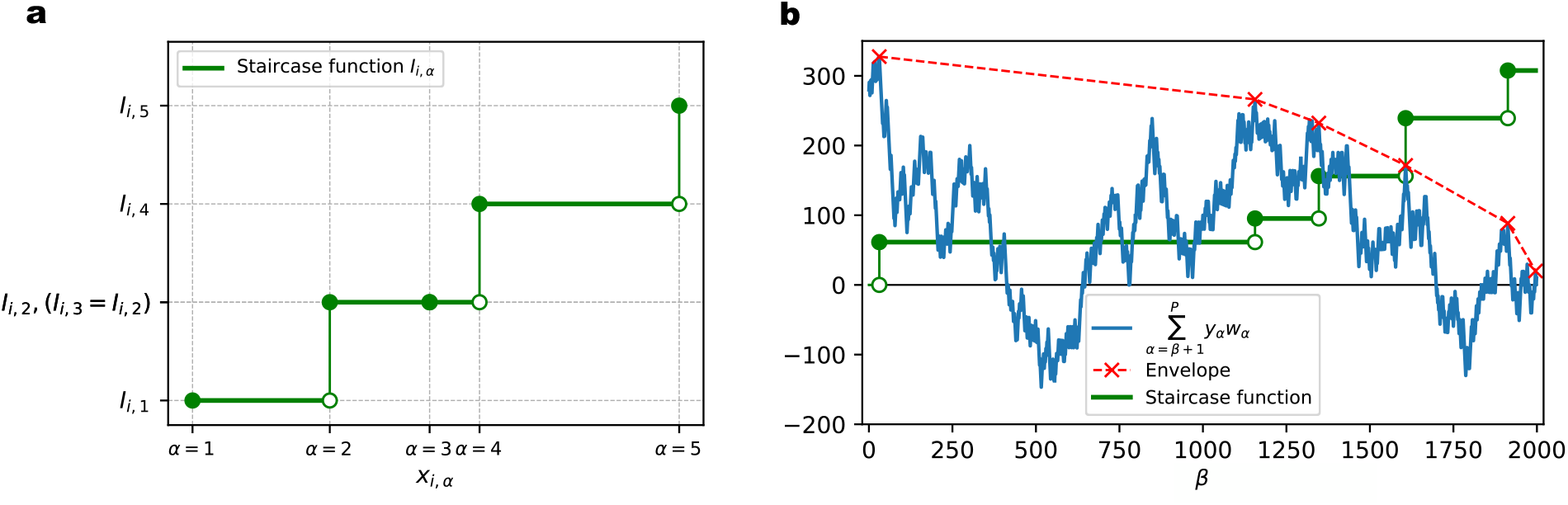
Illustration of the unrestricted model and its training algorithm. **(a)**. For the *i*-th axon, its aggregate transmission function value at input *x*_*i,α*_ is parameterized by the height of staircase function *I*_*i,α*_. The example shown here demonstrates 5 different input values and their corresponding aggregate synaptic function values on the *i*-th axon. The staircase function either increases (at *α* = 1, 2, 4 and 5) or stays flat (at *α* = 3) at each input value, in order to maintain monotonicity (nondecreasing). **(b)**. Envelope method to find the solution to eqns. (7) and (8), for each input dimension. Staircase functions (green solid line) are either flat or increase at different input values. Solid green dots indicate the increased function value. A (cumulative) random walk (blue solid line) is constructed from the current importance weights of the data points and their labels. An piece-wise linear envelope (red dashed line) is constructed such that it is always above or touching the (cumulative) random walk. The slopes of the envelope segments are decreasing from left to right.

As illustrated in Fig. 4c, networks with parallel synapses (*D*_hidden_ = 5, 10, 20 and 30, with 3 parallel synapses), show higher accuracy after 50 epochs of training compared to their counterparts with linear synapses and an equivalent number of parameters. Specifically, the improvement in accuracy ranges from 0.75% to 2.0%. This shows that multiple nonlinear parallel synapses can indeed improve the generalization performance on a supervised learning task with nontrivially structured data. The testing accuracy values for each network are listed in Supplementary Table 2. In Fig. 4d, we show the learned aggregate synaptic transmission functions of an example network, grouped by the output neuron to which they connect.

## Discussion

In this study, we have demonstrated that multiple nonlinear synapses between two neurons can enhance the memory capacity compared to a single linear connection. We have modeled the nonlinear transmission functions of each of these parallel synapses as a simple sigmoidal nonlinearity. Given different parameters (e.g. thresholds of the sigmoids), this allows the parallel synapses to contribute in a non-redundant fashion to the required computations, and thus may offer a functional explanation of the observation of multiple synaptic contacts between a pair of neurons in terms of enhanced classification capacity. The synaptic nonlinearity is essential here, since linear parallel synapses would be degenerate in the sense that their combined effect on the current into the postsynaptic neuron would still be linear.

Unlike in models incorporating nontrivial dendritic computation [29] or hidden layer neural networks, the nonlinearities in our model exclusively act on individual inputs. Consequently, our model’s architecture mirrors a Perceptron [31]. The sole nonlinearity that integrates multiple inputs and allows interactions between them is the step-like activation function at the soma, which implements the binary classification.

Specifically, we have systematically investigated the increase in memory capacity using random binary classification tasks, observing an increase of the capacity *P*^∗^*/N* with the number of input axons. We showed that even a small number of parallel synapses between two neurons leads to much larger capacity than in the classical Perceptron model. Furthermore, we examined a model with an a priori unlimited number of nonlinear synapses between two neurons. In this case the growth of the number of stored memory patterns with the number of input axons showed a superlinear behavior (similar to the previous model with sufficiently many parallel synapses). However, we found that only a relatively small number of potential synapses were utilized. The effective number of parallel synapses fell within the range observed in the cortex [4, 5, 9, 10, 21], suggesting that in the presence of a fixed number of available input axons, neurons in the brain may form parallel synapses until diminishing returns limit their capacity improvement. We also evaluated the generalization ability of a neural network equipped with parallel synapses. Using the MNIST dataset, we showed that parallel synapses enhance the test classification accuracy compared to networks with linear synapses, even when both models have the same number of parameters.

A fundamental assumption in our model is the nonlinearity and learnability of the synaptic transmission function. Synaptic transmission mechanisms involving stochastic vesicle release, the resulting resource depletion, and short-term synaptic plasticity are unlikely to result in postsynaptic currents that grow exactly linearly with the presynaptic firing rate. However, relating our models to these mechanisms will require a more biophysically detailed implementation involving spiking neurons. Also, while certain synaptic parameters such as the physical size of synapses, which is often considered as a proxy for the overall synaptic efficacy, can certainly change as a consequence of long-term potentiation or depression events, it is less clear to what extent more subtle synaptic properties, corresponding to the detailed shape of our synaptic transmission functions, can be reliably modified during biological learning.

Biological synapses on a given presynaptic neuron are typically either all excitatory or all inhibitory, which is known as Dale’s law [34]. They usually do not switch between excitation and inhibition, although evidence of neurotransmitter switching has been observed [35]. Previous research has considered the effects of sign-constrained synaptic weights on Perceptron learning [36] and classification capacity [37, 38]. In the learning algorithms for our models, such a sign constraint is already implemented. For example, in the restricted model, the aggregate synaptic function is a sum of several monotonically increasing sigmoid functions, and thus also monotonically increasing. For the unrestricted model, the aggregate synaptic function is parameterized as a sum of step-functions with nonnegative stepsizes, and thus is also nondecreasing. Therefore it would perhaps be appropriate to compare our models to the sign-constrained Perceptron, and since its capacity is only half as large as for the standard Perceptron [37], such a comparison would be even more favorable for our models.

This study has focused exclusively on offline learning, in which the algorithm has access to the whole dataset at any point during the learning process. However, a more biologically relevant situation would restrict the learning to occur one data point at a time. Adapting our algorithms to such an online learning scenario presents an interesting future direction for our research.

## Methods

### Binary classification task

In order to numerically quantify the memory capacity of our model, we utilize binary classification tasks of randomly generated input activity patterns. The binary labels associated with each pattern are also randomized. To generate an *N* -dimensional pattern **x**^*µ*^, we sample the input of the *i*-th axon (*i* = 1, 2, …, *N*) from a uniform distribution between 0 and 1, represented by *U* (0, 1), independently for each input dimension. The label *y*^*µ*^ associated with the pattern is also independently drawn from a Bernoulli distribution with equal probabilities for +1/ − 1.

### Estimating the capacity from numerical simulations

To evaluate the memory capacity of our model, we adopt a methodology similar to that used to measure the memory capacity of the Perceptron. We train a model neuron to classify random patterns. When the number of random patterns, denoted as *P*, is relatively small, the model neuron is likely to achieve successful classification. However, as the number of patterns increases, at some point the model neuron can no longer classify all the patterns correctly. We define a critical number of patterns, denoted as *P*^∗^, at which the model neuron has a 50% chance of successfully classifying all the patterns at the end of training.

The capacity (*P*^∗^*/N*) of the restricted model neuron depends on the number of input axons (*N*) and the number of parallel synapses per axons (*M*). We test various *P* values to assess the critical number of pattern (*P*^∗^) for a restricted model neuron. For each problem size *P*, we repeat the numerical experiments at least 5 times to estimate the successful classification rate for the model neuron (Fig. 2c-e). The capacity (*P*^∗^*/N*) is defined as the critical problem size *P*^∗^ normalized by the number of input axons. To find *P*^∗^*/N*, we need to interpolate the success rate between different values of *P/N*, which is achieved by logistic regression. Fitting a single logistic regression to the success rates obtained for a model neuron yields a point estimate of its capacity *P*^∗^*/N*. Additionally, we estimate the confidence interval of the capacity *P*^∗^*/N* following nonparametric bootstrap. We sample with replacement from the trials for each specific *M* and *N*. For each resampling, we fit a logistic regression to the successful classification rates as a function of *P/N*. Interpolating at a success rate of 0.5 with the fitted logistic regression model allows us to estimate a capacity value for that resampling. For each unique (*N, M*) configuration of the restricted model neuron, repeating this resampling process 100 times provides an estimation of the capacity and its confidence interval. This whole procedure is performed for every restricted model neuron with distinct (*N, M*) values.

For the unrestricted model neuron, the estimation method for the capacity is similar to that of the restricted model neuron. The only difference is that the capacity depends solely on the number of input axons (*N*). Therefore, we only need to determine the capacity for various *N* values. As above, the resampling technique is employed to estimate the confidence interval of the capacity, for each unrestricted model neuron with distinct *N* value.

### Restricted neuron trained with gradient descent

#### Model

In this version of the model neuron, the transmission function of each individual synapse is characterized by a sigmoid function (eqn. (1)). The *j*-th (*j* = 1 … *M*) parallel synapse on the *i*-th (*i* = 1 … *N*) axon has three parameters. The first parameter is *a*_*i,j*_. We take its squared value, (*a*_*i,j*_)^2^, as the amplitude of the transmission function, i.e., the maximum value of the response. The second parameter, *t*_*i,j*_, is the threshold of the transmission function, which represents the value of the presynaptic input at which the postsynaptic response increases most rapidly. The third parameter, *s*_*i,j*_, is the slope of the transmission function, which characterizes the sensitivity to input changes near the threshold. Since, in general, the synaptic response for excitatory synapses increases with larger input, we constrain the synaptic transmission function to increase monotonically (positive slope). That is, *s*_*i,j*_ is constrained to be nonnegative. The input for the *µ*-th pattern on the *i*-th axon is denoted by *x*_*i,µ*_.

Unlike models with nontrivial dendritic computation or hidden layer neural networks, the synaptic non-linearities in our model only act on individual input channels, and thus the model architecture is essentially that of a simple Perceptron. The only nonlinearity that combines multiple inputs and allows them to interact is the step-like activation function at the soma that implements the binary classification (or Softplus somatic nonlinearity for the MNIST simulations).

For each input **x**_*µ*_, the total current into the soma is 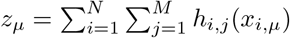. The prediction for *µ*-th data point is ŷ_*µ*_ = sign (*z*_*µ*_ − *θ*), where *θ* is a threshold parameter regulating the excitability of the neuron. This parameter can also be trained with gradient descent.

During simulations, we are using the hyperbolic tangent instead of the (positive) sigmoid function to model the individual synaptic transmission functions *h*_*i,j*_. Since tanh(*x*) is centered around 0, this makes it slightly easier to learn the final threshold *θ*, but otherwise this approach is mathematically equivalent, since the two functions are related as tanh(*x*) = 2*σ*(2*x*) − 1. Denote the parameters of hyperbolic tangent function as 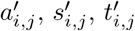 and *θ*^′^. The hyperbolic tangent function that we use in simulations is then

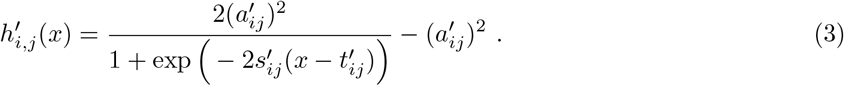

Comparing eqn. (1) and eqn. (3), the learned parameters using hyperbolic tangent and sigmoid function can be linked using the following equations: 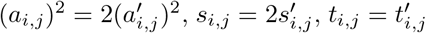 and 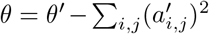 In Fig. 2h-j, we show the parameters of the sigmoid function used in the main text, which are essentially scaled versions of parameters in the hyperbolic tangent function.

#### Training

For the binary classification task, we use the hinge loss as the cost function (eqn. (4)). The parameter *ϵ* is the margin of the hinge loss, which we set to 0.1 during training:

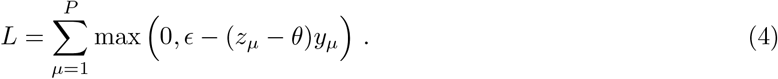

Based on gradient descent, the update rules of the parameters are as follows (eqn. (5)):

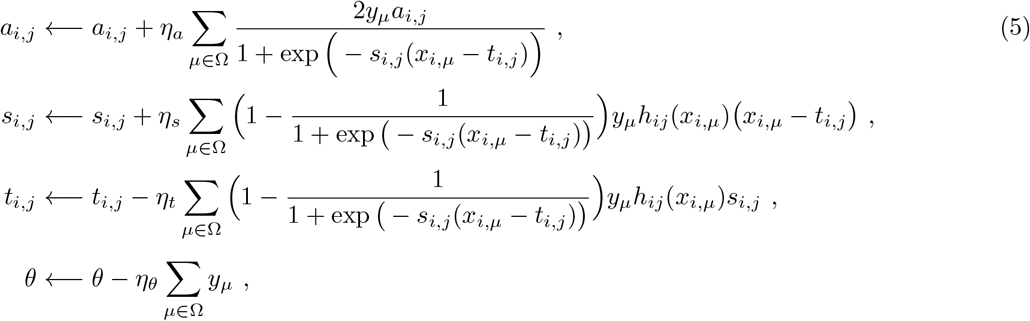

where the set Ω includes all data points that are misclassified before the update, or correctly classified by a margin less than *ϵ*, i.e., Ω : {*µ*|*ϵ* − (*z*_*µ*_ − *θ*)*y*_*µ*_ > 0}.

During training, we observe that the amplitude of certain synapses will become very small, which leads to a vanishing gradient of all parameters of these synapses, according to eqn. (5). Those synapses with extremely small amplitude will become ineffective, and their parameters will no longer be updated. To utilize those ineffective synapses again during training, we increase their amplitude (*a*_*ij*_)^2^ to a minimum value of 0.01. Their threshold parameters are also randomly shuffled to different values. The motivation is that the thresholds of those ineffective synapses may not be helpful for learning (causing their amplitude to become small), so randomly changing their thresholds to different values might increase the chance of better classification. The values of the shuffled thresholds are drawn from a distribution with support on the input range. This distribution has higher density near the edges of the input range, which mimics the learned threshold distribution (Fig. 2i).

### Training of the unrestricted neuron model

To parameterize the arbitrary monotonic transmission function from each axon of the unrestricted model, we assume that it takes the form of a staircase composed of step-like functions of the input value (Fig. 5). The model neuron is parameterized by *N* aggregate transmission functions in total, each for one axon. There are *P* data points in the dataset, thus there will be at most *P* different values the arbitrary monotonic function can take on this dataset for each input dimension. In other words, we assume that there are up to *P* (step-like) parallel synapses on each axon, which limits the biological plausibility of this model.

We constrain each aggregate synaptic function to be monotonic, i.e, for excitatory synapses the function values have to be increasing with increasing input, for each input dimension. We label the synaptic function value of the *α*-th input value on the *i*-th axon as *I*_*i,α*_, where the *α* index labels the data points in order of increasing input values (for that axon).

To correctly classify all data points, our training algorithm has to construct a suitable monotonic function, *I*_*i,α*_, for each axon. For the *µ*-th data point, the input is *x*_*i,µ*_ on *i*-th axon. If its label *y*_*µ*_ is +1, the algorithm tends to increase the value at *x*_*i,µ*_, whereas if *y*_*µ*_ is −1, the algorithm pushes *I*_*i,µ*_ towards the negative direction. Note that there are two types of ordering of the data points involved. The first one is the natural ordering of the data points, which is indexed by *µ* and shared by all axons. The other one is the order of increasing inputs *x*_*i,µ*_, which is indexed by *α*. Since the increasing order of *x*_*i,µ*_ differs for each dimension *i*, this ordering described by *α* depends on *i*. The two types of ordering can be linked by a permutation matrix for each axon, which is completely determined by the input data. We will use *x*_*i,α*_ (with input values increasing with *α*) in the following for simplicity.

We can use − Σ_*α*_ *y*_*α*_*I*_*i,α*_ as part of our cost function for the *i*-th axon, which will push the aggregate synaptic function in the correct direction corresponding to the label *y*_*α*_ of each data point with a given input value *x*_*i,α*_. Note that the computation of *I*_*i,α*_ can be performed in parallel for different input dimensions *i*. Therefore, we drop the index of *i* in the remainder of this section for simplicity. Since there could be data points that are more difficult to classify than other data points, we introduce an importance weight *w*_*α*_ for each data point in the loss function by writing − Σ_*α*_ *y*_*α*_*w*_*α*_*I*_*α*_. This leads to an efficient, iterative way to calculate *w*_*α*_, such that more difficult data points can have higher importance weights in the next iteration of learning.

The complete cost function we employ for each input dimension is 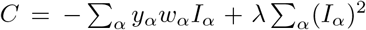. Here Σ_*α*_(*I*_*α*_)^2^ is an L2 regularization term with coefficient *λ* that discourages postsynaptic currents from becoming too large. To satisfy the monotonicity constraint of the aggregate transmission function, it has to obey *I*_1_ ≤ *I*_2_ ≤ · · · ≤ *I*_*P*_. To achieve this, we introduce the step size parameters *ρ*_*β*_, with *β* = 1 … *P* − 1, such that 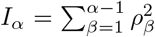, where without loss of generality we have imposed *I*_1_ = 0. Here 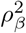 represents the step size of the staircase function at the *β* + 1-th input value. Then we formulate an optimization problem as follows

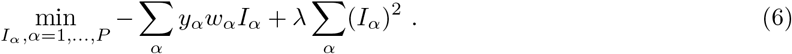

Taking the first and second derivative of *C* = − Σ_*α*_ *y*_*α*_*w*_*α*_*I*_*α*_ + *λ Σ*_*α*_(*I*_*α*_)^2^ with respect to *ρ*_*β*_, we find

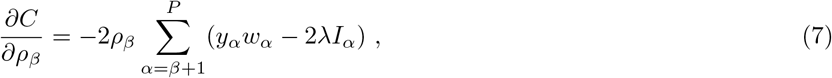

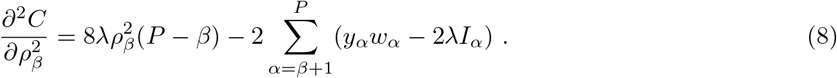

Setting the first derivative to zero, *ρ*_*β*_ and *I*_*α*_ have to satisfy either *ρ*_*β*_ = 0 or 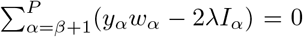 for each *β*. The case *ρ*_*β*_ = 0 means that the step function is flat at *β* + 1-th input value (no step). The alternative case 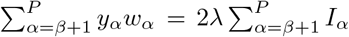 means that the function increases at the *β* + 1-th input value. However, to keep 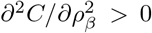, the two conditions cannot be satisfied at the same time, so our algorithm has to decide which one should hold for each *β*.

Solutions to the optimization problem specified by eqn. (7) and eqn. (8) can be found by using the envelope method illustrated in Fig. 5. The problem is equivalent to determining values of all *I*_*β*+1_ where the step-sizes are nonzero (such that *I*_*β*+1_ *> I*_*β*_). The envelope method iteratively determines *I*_*β*+1_ with nonzero staircase step size from right to left (*β* = *P* to *β* = 1). Suppose *β* + 1 is the rightmost location (*β* is closest to *P*) where the staircase step size is nonzero, i.e., *I*_*α*_ = *I*_*β*+1_ for all *α* ≥ *β* + 1, then from eqn. (7), we have

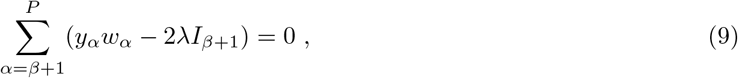

and thus

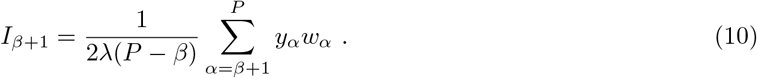

Based on eqn. (10), the value of *I*_*β*+1_ can be determined using all the *y*_*α*_ and *w*_*α*_ to its right (for *α* ≥ *β* + 1). If we consider *y*_*α*_*w*_*α*_ as the step of random walk at each *α* (depending on the random labels *y*_*α*_ and the importance weights), then 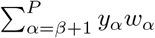 is the cumulative distance traversed between *β* + 1 and *P*.

For the remaining locations with nonzero staircase step sizes at smaller *β* (corresponding to data points with smaller input values), we can use eqn. (7) and eqn. (8) in a similar fashion. Suppose *β*_1_ + 1 and *β*_2_ + 1 are two adjacent locations where the staircase step sizes are nonvanishing (with *β*_2_ > *β*_1_), then we have

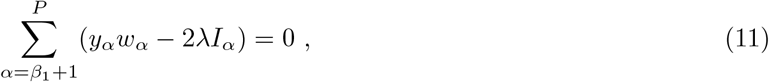

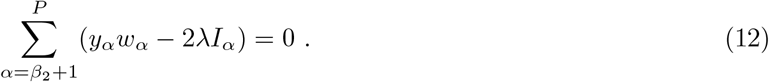

Subtracting eqn. (12) from eqn. (11), we obtain

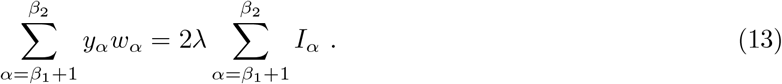

Since *β*_1_ and *β*_2_ are two adjacent steps, all *I*_*α*_ between them (with *α* = *β*_1_ + 1, …, *β*_2_) have the same value, i.e.,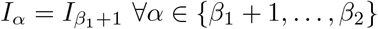. This leads to

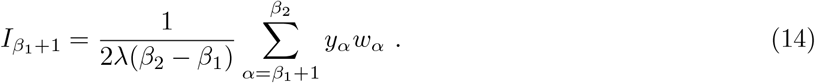

As above, if considering *y*_*α*_*w*_*α*_ as the step of a one-dimensional random walk at each *α*, the quantity 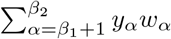 is the cumulative distance traveled between *β*_1_ and *β*_2_. The intuition that follows from eqn. (10) and eqn. (14) leads us to construct an envelope of the random walk, as in Fig. 5, such that the slopes of each segment are negative. The steps sizes of our staircase function are only nonzero at locations where the envelope touches the random walk. The value of the staircase function is proportional to the slope of the envelope. Since the synaptic functions are monotonic (nondecreasing in this case), the slopes of the envelope have to be nonincreasing as *α* grows.

Having solved the above optimization problem separately for each axon, we can now tackle the overall classification problem using the following iterative process: 1. Given the ordered input data *x*_*i,α*_, labels *y*_*α*_ and the current importance weights *w*_*α*_ (whose values are all initialized to unity), construct the envelope and calculate the aggregate synaptic transmission function *I*_*α*_; 2. Given the new *I*_*α*_, evaluate the model’s prediction for all the data points; 3. Re-weight the data points by updating *w*_*α*_ if the data point *α* is misclassified, incrementing *w*_*α*_ ← *w*_*α*_ + 1. We repeat steps 1 to 3, until all data points are correctly classified or a maximum iteration number is reached.

### Training neural network models with parallel synapses on the MNIST classification task

For the MNIST classification task, we employ a fully connected neural network with one hidden layer. This network processes images of handwritten digits from the MNIST dataset, each being 28 × 28 pixels. The network’s input layer consists of 784 neurons, corresponding to the pixel count, and the output layer has 10 neurons, representing a one-hot encoding of digit identities ranging from 0 to 9. Linear (single) synapses form the feedforward connections from the input layer to the hidden layer. The hidden layer’s nonlinearity is a Softplus function, which is a smooth approximation to the Rectified Linear (ReLU) function. The feedforward connections from the hidden layer to the output layer are made through either nonlinear parallel synapses or single linear synapses (for comparison). In both types of networks, we apply batch normalization of inputs to the hidden layer. We employ a multi-label cross-entropy loss for training. The hyperparamter settings are described in the Supplementary Table 3.

In networks with parallel synapses, these synapses are used exclusively from the hidden layer to the output layer. Each set of parallel synapses, connecting a pair of neurons, is constrained to have a monotonically increasing aggregate transmission function. For the purpose of fair comparison, the linear synapses connecting hidden layer and output layer in networks without parallel synapse are also constrained to be monotonically increasing, i.e., have positive weights.

The MNIST dataset [27] comprises 60,000 images in the training set and 10,000 images in the testing set. During each training epoch, we train the networks on all images in the training set and subsequently test their classification accuracy on the testing set.

## Code and data availability

Code to reproduce these results will be made available on GitHub.

## Acknowledgements

The numerical simulation were performed on the Nautilus platform, which is supported in part by National Science Foundation (NSF) awards CNS-1730158, ACI-1540112, ACI-1541349, OAC-1826967, OAC-2112167, CNS-2100237, CNS-2120019, the University of California Office of the President, and the University of California San Diego’s California Institute for Telecommunications and Information Technology/Qualcomm Institute. Thanks to CENIC for the 100Gbps networks. M.K.B was supported by R01NS125298 (NINDS) and the Kavli Institute for Brain and Mind.

## Supplementary Materials

### Comparison of two types of neural networks for MNIST task

Here we enumerate the total number of parameters in networks with nonlinear parallel synapses and networks with standard linear synapses. We use *D*_in_ as the number of neurons in the input layer, which is 28 ∗ 28 for both types of network. *D*_out_ denotes the number of neurons in the output layer, which is 10 for both types of network. *D*_hidden_ is the number of neurons in the hidden layer. We choose *D*_hidden_ as 5, 10, 20 and 30 for networks with parallel synapses. We set the number of parallel synapses per connection to 3, i.e., *M* = 3. For networks with nonlinear parallel synapse in the hidden-output connection, the total number of parameters is (*D*_in_ + 1)*D*_hidden_ + (3*MD*_hidden_ + 1)*D*_out_, including the bias terms. For networks with only single linear synapses, the total number of parameters is (*D*_in_ + 1)*D*_hidden_ +(*D*_hidden_ + 1)*D*_out_, also including the bias terms. Therefore, we opt for a slightly larger *D*_hidden_ for networks with linear synapses to achieve a fair comparison. The parameter counts for the two types of networks are shown below in Table 1.

**Table 1:**
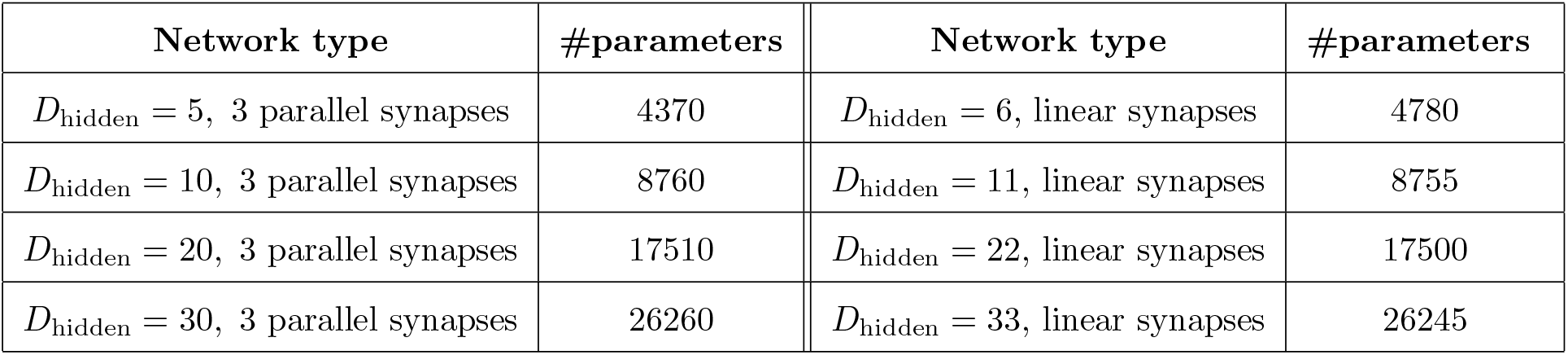
Comparison of parameter numbers in networks with parallel synapses (left two columns) and networks with single linear synapses (right two columns).

Below we record the accuracy of both types of networks on the testing set after 50 epochs of training in Table 2. The standard deviation is calculated from 20 trained networks with different initialization seeds.

**Table 2:** Comparison of the classification accuracy in networks with parallel synapses (left two columns) and networks with single linear synapses (middle two columns). The accuracy improvements from using parallel synapses are shown in the right column.

| Network type | Accuracy | Network type | Accuracy | Gain |
| --- | --- | --- | --- | --- |
| $D_{\text{hidden}} = 5$, 3 parallel synapses | $91.42\% \pm 0.06\%$ | $D_{\text{hidden}} = 6$, linear synapses | $89.93\% \pm 0.11\%$ | **1.49%** |
| $D_{\text{hidden}} = 10$, 3 parallel synapses | $94.92\% \pm 0.06\%$ | $D_{\text{hidden}} = 11$, linear synapses | $92.93\% \pm 0.04\%$ | **2.00%** |
| $D_{\text{hidden}} = 20$, 3 parallel synapses | $96.41\% \pm 0.05\%$ | $D_{\text{hidden}} = 22$, linear synapses | $94.90\% \pm 0.03\%$ | **1.51%** |
| $D_{\text{hidden}} = 30$, 3 parallel synapses | $96.77\% \pm 0.03\%$ | $D_{\text{hidden}} = 33$, linear synapses | $96.02\% \pm 0.02\%$ | **0.75%** |

**Table 3:**
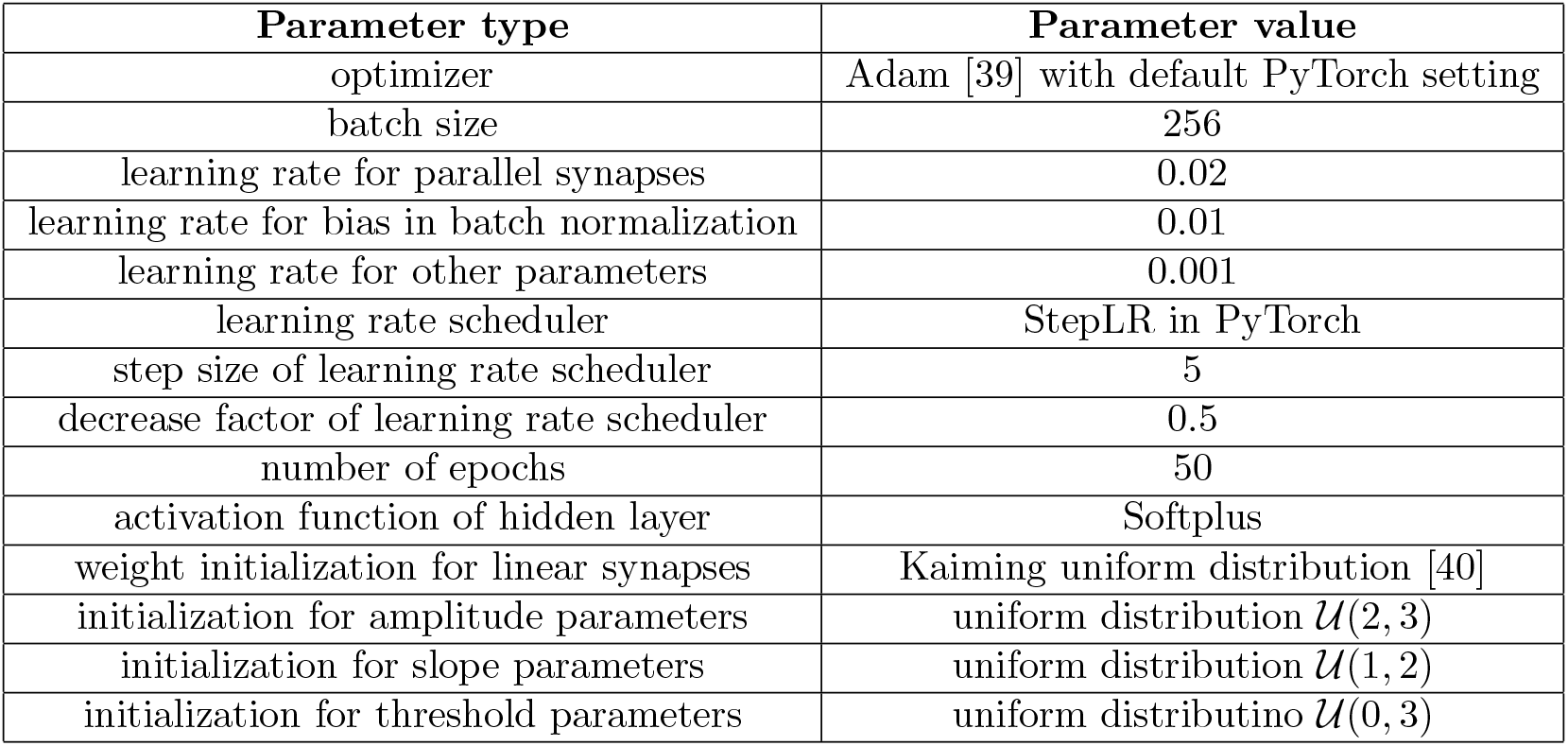
Hyperparameter settings for training neural networks.

### Learned aggregate synaptic function in neural networks

For the network depicted in Fig. 4a (with *D*_hidden_ = 10, 3 parallel synapses), we can also visualize the activation distributions of individual hidden units, as shown in Fig. 6. The input patterns are from the testing set of the MNIST dataset. For each hidden unit, we collect its activations (before the nonlinear synaptic transmission function) across various input patterns. These activations serve as the input values to the parallel synapses. Each hidden unit is connected to each output unit via a set of parallel synapses. A set of parallel synapses can be represented by their aggregate synaptic function. Thus, for each hidden unit, we can plot all the aggregate synaptic functions connecting this hidden unit to all the output units, which is also visualized in Fig. 6. In addition, Fig. 7 shows the distributions of parameters (slope, amplitude and threshold) in the parallel synapses from the same network.

**Figure 6:**
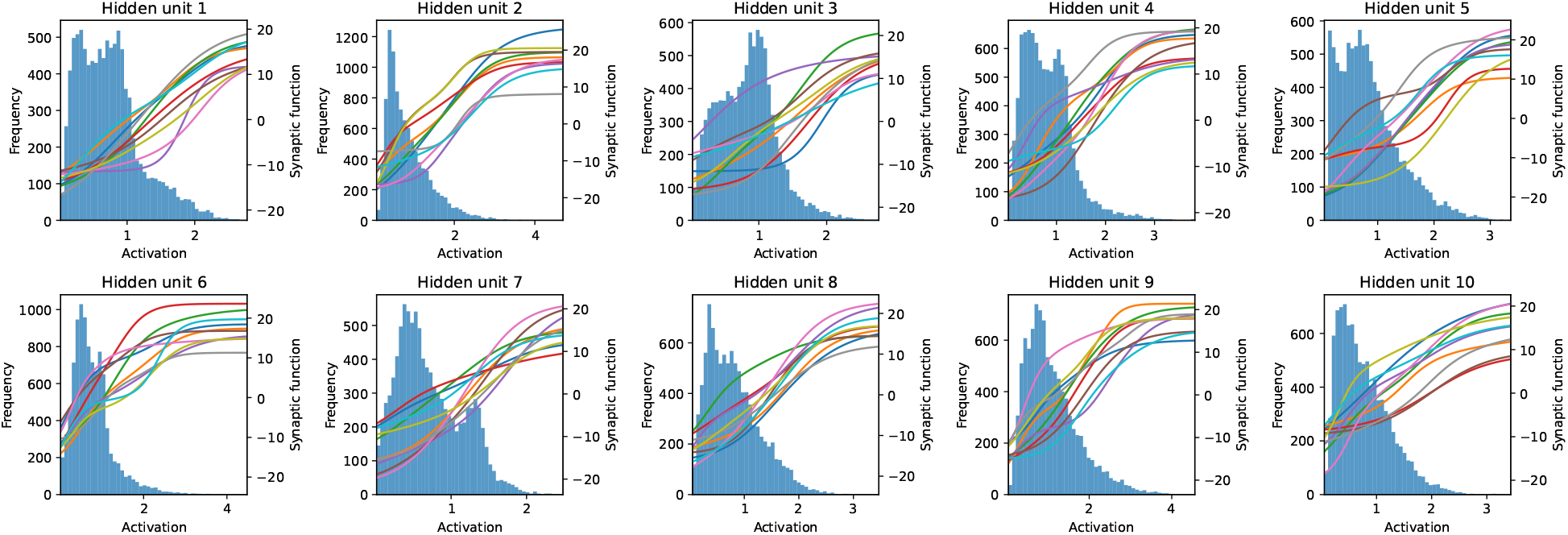
Histograms of hidden unit activations in the two-layer neural network with parallel synapses, overlaid with learned aggregate synaptic transmission functions. The x-axis is the total activation of the corresponding hidden unit. The left y-axis is the frequency of activation values. The right y-axis is the value of the aggregate synaptic function. Each panel corresponds to one hidden unit. The histograms collect activation values from all patterns in the testing set of the MNIST dataset. The lines are aggregate synaptic functions connecting each hidden unit to all output units, with 10 output units in total.

**Figure 7:**
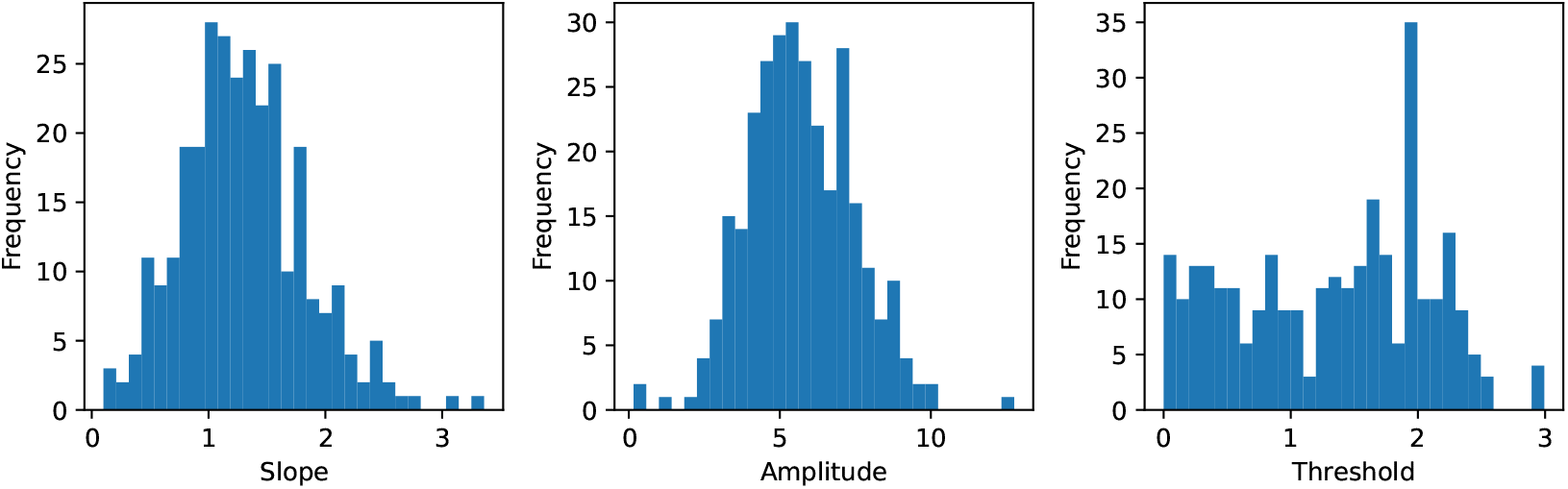
Histograms of parameters in parallel synapses from the same network as in Fig. 4, with slope (left), amplitude (middle) and threshold (right).

## Notes

### Competing Interest Statement

The authors have declared no competing interest.

